# Integrated Genomic Profiling Reveals Mechanisms of Broad Drug Resistance and Opportunities for Phenotypic Reprogramming in Cancer Cells

**DOI:** 10.1101/2025.09.04.674202

**Authors:** Ian Mersich, Brian S. J. Blagg, Aktar Ali

## Abstract

Broad drug resistance is a major barrier to effective cancer therapy, driven by diverse genetic, transcriptional, and metabolic adaptations across tumor types. Here, we developed an integrative computational framework that leverages PRISM drug sensitivity profiles from DepMap, multi-omic datasets, and perturbagen libraries to systematically characterize and identify strategies to reverse broad resistance in cancer cell lines. We found that resistant lines exhibit transcriptional programs enriched for extracellular matrix remodeling, stress adaptation, and survival signaling, with NFE2L2 emerging as a central regulatory hub linked to upstream mutations and downstream oxidative stress pathways. Integrated metabolomics and transcriptomics highlighted metabolic reprogramming as a hallmark of resistance, while mutation analyses revealed convergence on growth factor and ECM-related pathways. These features were also reflected in patient cohorts, where resistance-associated mutations correlated with reduced progression-free survival across diverse cancer types. Computational perturbagen screening identified candidate compounds predicted to reverse resistance-associated gene expression profiles, converging on actionable targets including NFE2L2, ABCB1, and CYP3A4, with compounds such as brefeldin A and nocodazole predicted to have strong activity in resistant lines. This study establishes a scalable, mechanism-informed framework for rationally identifying and prioritizing compounds to overcome broad drug resistance in cancer, providing a roadmap for targeted re-sensitization strategies.

## INTRODUCTION

Drug resistance remains a central barrier to effective cancer treatment, arising through diverse and often multifactorial mechanisms that allow cancer cells to evade the cytotoxic effects of therapy^1-3^. Mechanisms driving resistance include enhanced drug efflux via transporters^4-7^, alterations in drug targets^8^, dysregulation of apoptosis pathways^9-11^, and activation of stress-response signaling networks^12-14^. Additionally, cancer stem-like cells, cellular quiescence, epithelial-to-mesenchymal transition, and heterogenous cell populations have been implicated in mediating broad and persistent resistance phenotypes across cancer types^15-19^. Understanding and overcoming these resistance mechanisms is essential for advancing durable cancer therapies and improving patient outcomes^20,21^.

The continued emergence of new mechanisms of multidrug resistance, including adaptations in metabolic pathways and microenvironmental interactions, further underscores the complexity of drug resistance in cancer and the need for integrative strategies to address it^22^. Advances in large-scale functional genomics and pharmacologic screening efforts, such as the Cancer Dependency Map^23^ and PRISM Repurposing datasets^24^, have enabled systematic mapping of drug responses across thousands of genetically characterized cancer cell lines, providing a framework to investigate the molecular correlates of drug sensitivity and resistance at unprecedented scale^25-29^. These resources facilitate integrated approaches that combine drug response profiling with multi-omic data, including transcriptomics, mutational landscapes, and metabolomics, to identify pathways and gene networks that mediate resistance phenotypes^30-33^. The availability of comprehensive clinical and genomic resources, such as the Catalogue of Somatic Mutations in Cancer (COSMIC)^34^, The Cancer Genome Atlas (TCGA)^35^ and ClinVar^36^, alongside bioinformatics toolkits for drug resistance analysis^37^, further expands the capacity to systematically interrogate the genetic basis of therapeutic failure and prioritize candidate vulnerabilities for intervention.

Despite these advances, overcoming broad and multi-class drug resistance across mechanistically diverse therapies remains a formidable challenge in oncology. Efforts to systematically map resistance mechanisms and identify re-sensitization strategies are critical for the development of rational drug combinations and targeted interventions that can restore sensitivity in resistant cancers^24,38-45^. Here, we leverage large-scale drug sensitivity datasets in conjunction with transcriptomic, mutational, and metabolomic data to characterize the molecular features of broad drug resistance in cancer cell lines and to identify candidate compounds predicted to reverse resistant phenotypes through integrated perturbagen screening. By combining computational network analyses with pharmacogenomic data, this study aims to advance rational strategies for overcoming drug resistance and guiding the development of targeted therapies in resistant cancer models.

## RESULTS

### Systematic identification of broadly sensitive and resistant cancer cell lines

To explore shared mechanisms of broad-spectrum drug resistance, we developed a computational framework to classify cancer cell lines based on their global drug sensitivity profiles. The experimental workflow is outlined in Figure 1A-C. Using PRISM Repurposing data from the DepMap portal we computed a drug sensitivity score for each cell line by calculating the median log2 fold change for all compounds using subsets of mechanistically distinct drug classes (e.g., compounds with unique mechanism of action, kinase inhibitors, DNA-damaging agents, and epigenetic modulators), and averaged the class-level medians to generate a single composite sensitivity score (Figure 1A; Supplemental Table S.1). These scores were visualized against the overall drug response distribution across 6765 compounds (median log2 fold change for all compounds) revealing a broad range of sensitivities across cell lines. Transcriptomic (RNA-seq) and mutation profiles from DepMap were integrated to identify resistance-associated features, and enriched gene sets, linking mutations to differentially expressed genes (DEGs), were interrogated using network-level approaches (Figure 1B). In parallel, DEGs from resistant lines were queried against the LINCS L1000 database to identify perturbagens predicted to reverse resistance-associated expression signatures (Figure 1C).

**Figure 1.**
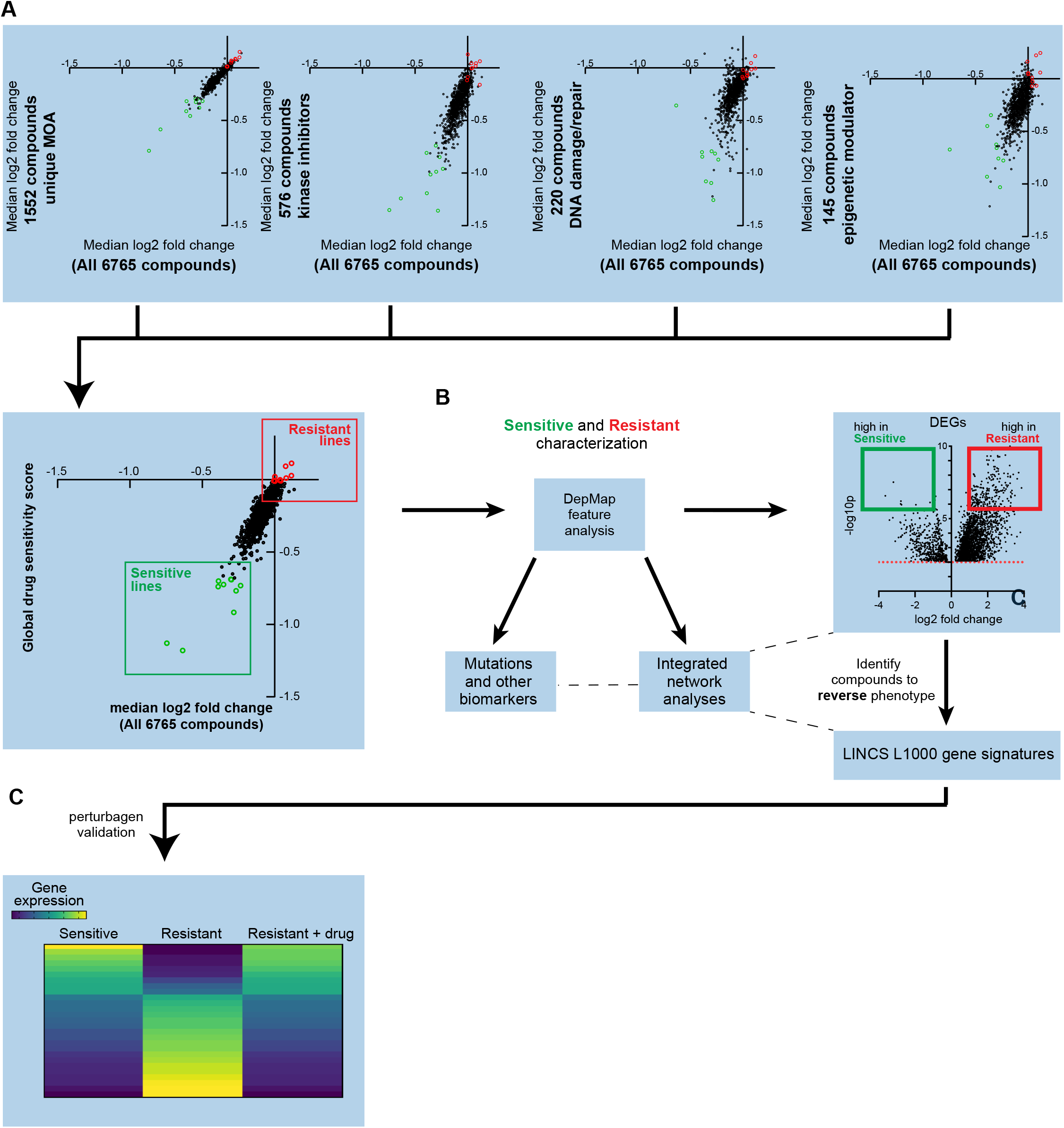
Systematic identification of broadly sensitive and resistant cancer cell lines. (A) Schematic of drug sensitivity scoring using median log2 fold change of PRISM repurposing compounds across mechanistically distinct drug classes. (B) Workflow for classifying cell lines as sensitive or resistant and comparing genomic features to characterize these populations. (C) Strategy for utilizing resistance-associated DEGs with perturbagen screening data to identify candidate compounds predicted to reverse the drug-resistant phenotype.

### Transcriptomic analysis reveals gene signatures and pathways associated with broad resistance

To investigate transcriptional programs underlining broad drug resistance, we performed differential expression analyses between resistant and sensitive cell lines using three complementary comparison strategies (Figure 2A– C; Supplemental Table S.2). Group 1 was defined by selecting the top and bottom 10% of lines based on composite drug sensitivity scores (n=91 in each group; Figure 2A). Group 2 consisted of transcriptionally clustered lines from group 1 that tightly segregated with their respective phenotypes, thereby minimizing intra-group heterogeneity (Figure 2B). Group 3 was constructed to control for lineage-specific biases by including only cancer types represented in both resistant and sensitive categories. Although there was some lineage imbalance in Groups 1 and 2 (Supplemental Figure 2A), univariate regressions analysis did not reveal any statistically significant association between drug sensitivity score and lineage (Supplemental Figure 2B). Additional metadata variables such as collection site, disease subtype, and media conditions were also evaluated but deemed unsuitable as covariates due to weak or inconsistent effects (Supplemental Figures 2C–F; Supplemental Tables S.3 and S.4). All three groups showed robust stratification between sensitive and resistant populations (Supplemental Figures 2G–I).

**Figure 2.**
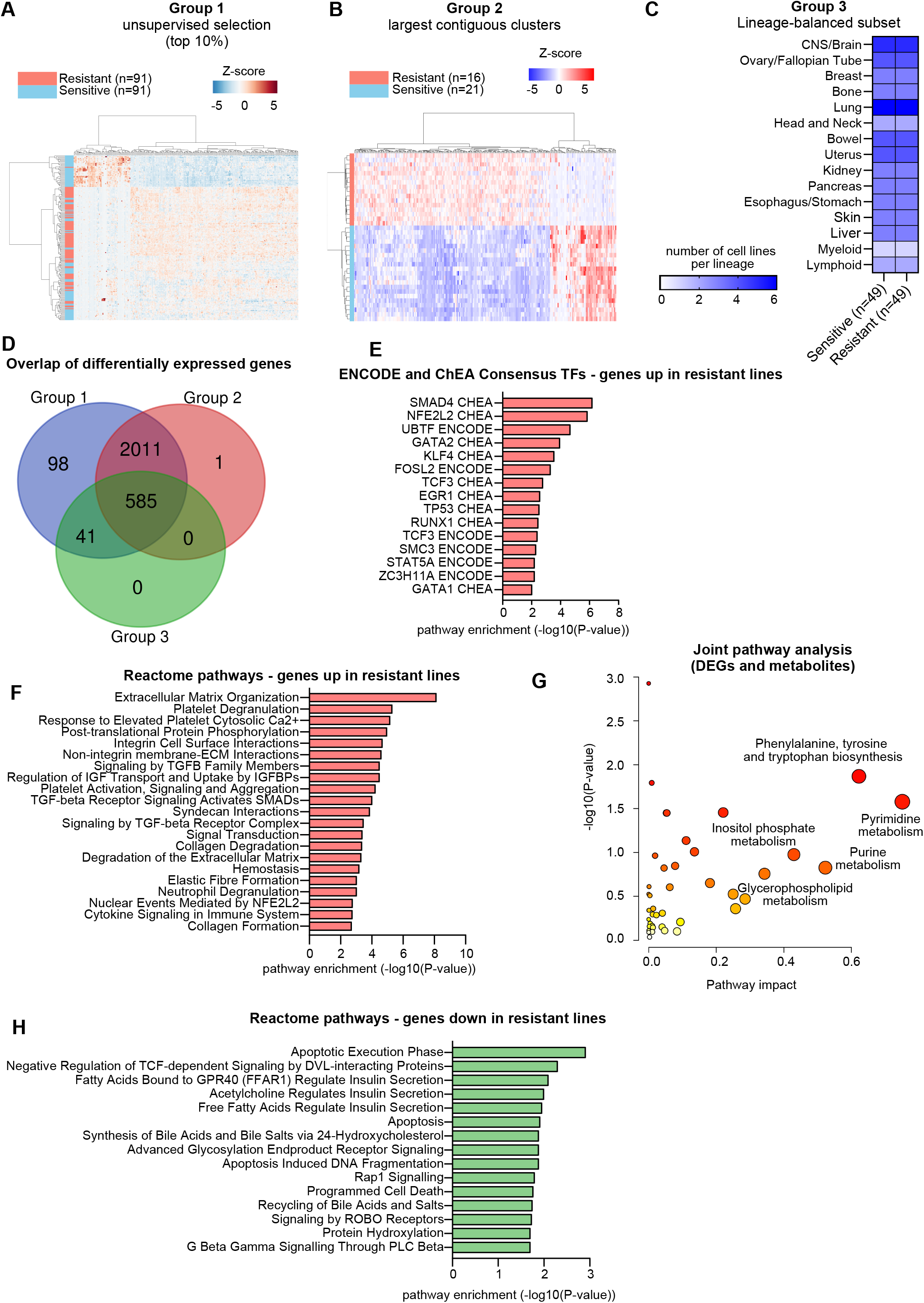
Transcriptomic analysis reveals gene signatures and pathways associated with broad resistance. (A–C) Schematic of three strategies for defining resistant and sensitive groups: Group 1, top and bottom 10% by drug sensitivity score; Group 2, largest contiguous transcriptional clusters (from Group 1); Group 3, lineage-balanced subsets. (D) Overlap of differentially expressed genes (DEGs) identified across the three group comparisons. (E) Enrichment analysis (Enrichr) of transcription factors regulating genes up in resistant lines. (F) Reactome pathway enrichment analysis (Enrichr) of genes upregulated in resistant lines. (G) Joint pathway analysis (Metaboanalyst) integrating DEGs and differentially abundant metabolites. (H) Reactome pathway enrichment analysis (Enrichr) of genes downregulated in resistant lines.

DEGs were identified for each group, and correlation analyses were conducted between gene expression and drug sensitivity scores (Supplemental Table S.5). A set of 585 DEGs (P<0.05) was shared across all three comparison groups, representing the most robust gene expression markers for broadly resistant cell lines (Figure 2D; Supplemental Table S.6).

Pathway enrichment of genes upregulated in resistant lines (P<0.05, effect size > 1) revealed several upstream transcription factors implicated in drug resistance, including NFE2L2 (NRF2), a central regulator of oxidative stress response and multidrug resistance, which ranked among the top hits in the ENCODE and ChEA Consensus TF gene set library (Figure 2E; Supplemental Table S.7). Reactome enrichment of these genes (Figure 2F; Supplemental Table S.8) identified several biologically relevant pathways associated with resistance, including extracellular matrix (ECM) organization, platelet activation, TGF-beta receptor signaling, IGF transport regulation, neutrophil degranulation, and Nuclear Events Mediated by NFE2L2. These results highlight a coordinated transcriptional program involving matrix remodeling, stress signaling, and survival mechanisms.

To complement the transcriptomic findings, we compared the metabolomics profiles between sensitive and resistant lines. Metabolites showing differential abundance were combined with DEGs in a joint pathway analysis using MetaboAnalyst (Figure 2G; Supplemental Table S.9 and 10). Several pathways emerged as significantly enriched, including amino acid biosynthesis (e.g., phenylalanine, tyrosine, valine, leucine), purine and pyrimidine metabolism, glycosaminoglycan degradation, and inositol phosphate metabolism. These results suggest that broad drug resistance is accompanied by widespread metabolic reprogramming affecting both nucleotide and amino acid metabolism.

Finally, Reactome pathway analysis of genes downregulated in resistant lines (Figure 2H) revealed enrichment for apoptosis-related programs, indicating that suppression of cell death machinery is a key feature of resistance. Taken together, these results suggest that broadly resistant cell lines undergo coordinated remodeling of stress adaptation, metabolic reprogramming and enhancement of survival programs.

### Mutation signatures and regulatory network analyses associated with broad drug resistance

To investigate mutation patterns associated with broad drug resistance, we applied two complementary analytical strategies. In the first approach, we compared gene-level mutation frequency differences between the 50 most resistant and 50 most sensitive cell lines (Figure 3A). For each gene, we calculated a “group skew score” as the difference in the percentage of mutated resistant lines versus sensitive lines. The top-ranked genes from each group were independently subjected to pathway enrichment analyses (Figures 3B and 3C).

**Figure 3.**
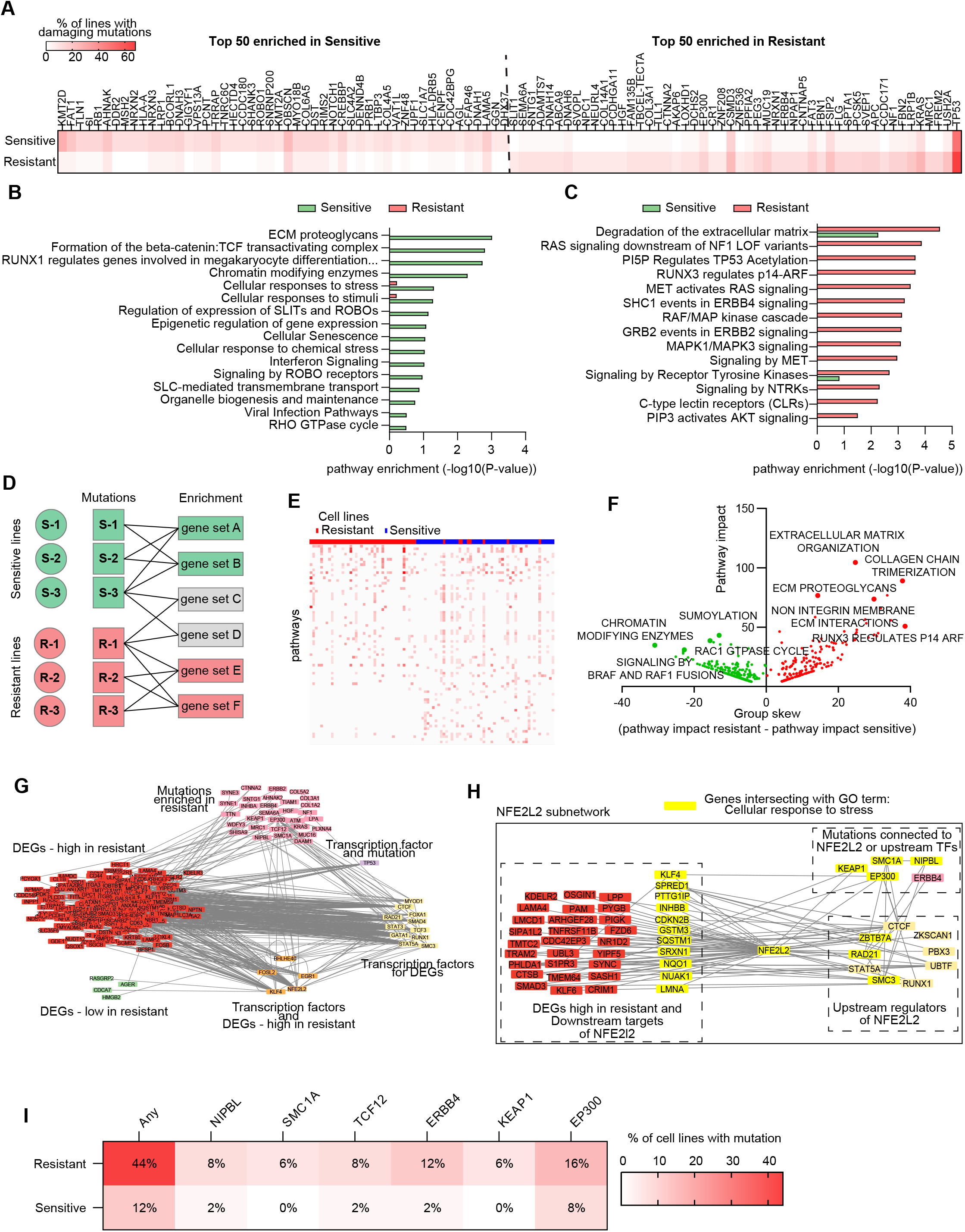
Mutation signatures and regulatory network analyses associated with broad drug resistance. (A) Gene-level mutation frequency differences between the 50 most resistant and 50 most sensitive cell lines. Top 50 genes ranked by group skew shown. (B-C) Pathway enrichment analyses of top genes preferentially mutated in sensitive (B) and resistant (C) lines. (D) Schematic of pathway-level mutation burden scoring per cell line. (E) Heatmap of pathway enrichment scores across resistant and sensitive lines. (F) Volcano plot summarizing pathway-level group skew and significance, highlighting pathways enriched in sensitive or resistant lines. (G) Integrative gene–regulator network combining DEGs, enriched transcription factors, and recurrently mutated genes. (H) NFE2L2-centered subnetwork highlighting upstream regulators and downstream targets linked to stress response and matrix remodeling. (I) Heatmap showing the percentage of resistant and sensitive lines with mutations in NFE2L2-linked genes, illustrating convergent and non-overlapping mutation patterns contributing to resistance.

Genes more frequently mutated in sensitive lines were enriched in pathways related to chromatin remodeling, cell cycle regulation, and cellular stress response, including chromatin modifying enzymes, RUNX1-regulated transcription, epigenetic regulation of gene expression, and cellular senescence, along with signaling cascades such as Rho GTPase signaling, ROBO receptor signaling, and interferon signaling (Figure 3B; Supplemental Table S.11). Interestingly, this gene set also implicated pathways tied to SARS-CoV-2 infection and viral host interactions, suggesting that heightened innate immune signaling or cellular stress priming may sensitize these lines to therapy.

Conversely, genes more commonly mutated in resistant lines were enriched for growth factor signaling, matrix remodeling, and canonical oncogenic pathways including RAF/MAPK signaling cascade, PI3K/AKT signaling, RTK signaling (e.g., MET, ERBB4, NTRK), and TP53 regulatory pathways (Figure 3C; Supplemental Table S.11). Notably, multiple pathways tied to extracellular matrix organization and immune system modulation were enriched, highlighting potential links between microenvironmental remodeling and broad drug resistance.

As a complementary strategy, we performed sample-specific pathway enrichment using the full mutation profiles of individual cell line (Figure 3D). Each sample was scored for pathway mutation burden based on -log10 p-values, and hierarchical clustering revealed distinct separation between resistant and sensitive lines (Figure 3E). To quantify these differences, we calculated pathway-level skew scores, defined as the difference in the proportion of enriched resistant versus sensitive samples, adjusted for the mean enrichment significance across each group. A volcano plot (Figure 3F; Supplemental Table S.12) highlighted key pathways skewed toward either phenotype. Sensitive lines exhibited enrichment for pathways such as second messenger signaling, WNT/TCF-dependent transcription, histone demethylation, and cell death regulation. Resistant lines showed enrichment in RUNX3/p14-ARF signaling, PDGF/MET/FGFR signaling, collagen biosynthesis, and non-integrin ECM interactions. Interestingly, Glycosaminoglycan degradation, a pathway involved in ECM turnover, was significantly enriched in resistant lines in both the mutation-based pathway analysis (Figure 3F) and prior joint transcriptomic-metabolomic pathway analysis (see also Supplemental Table S.10 from Figure 2G). Consistent with this, genes related to ECM proteoglycans were also enriched in the mutation analyses; however, the directionality of enrichment varied depending on the analytical strategy: gene-level mutation skew favored sensitive lines (see Figure 3B), while pathway-level mutation burden was enriched in resistant lines (see Figure 3F). This divergence highlights how analytical granularity, focusing on top-ranked genes versus full pathway in individual samples, can yield different biological interpretations, underscoring the complex and multifaceted role of ECM remodeling in drug response.

To further investigate the mechanistic connections between mutational patterns and resistance phenotypes, we constructed an integrative gene–regulator network (Figure 3G), incorporating frequently mutated genes, differentially expressed genes (DEGs), and transcription factors (TFs) enriched from ENCODE and ChEA databases. This network identified hubs of regulatory convergence, linking upstream mutations to downstream transcriptional changes. One prominent hub was NFE2L2, a key regulator of oxidative stress response and cell survival. A focused subnetwork centered on NFE2L2 (Figure 3H) revealed associations with xenobiotic metabolism, ECM remodeling, and antioxidant defenses. Several infrequently mutated genes (KEAP1, EP300, ERBB4, TCF12, SMC1A, and NIPBL) were functionally linked to this hub. Although each gene individually had a low mutation frequency of only (6–16%), 44% of resistant lines harbored at least one mutation in this set (Figure 3I). These mutations typically occurred in a mutually exclusive manner, supporting a model in which diverse genetic alterations converge to dysregulate NFE2L2 signaling, thereby promoting resistance across multiple drug classes.

### Clinical relevance of resistance-associated mutations in patient cohorts

To assess the clinical relevance of mutations identified in resistant cell lines, we queried cBioPortal using data from over 11,000 patients across 32 TCGA PanCancer Atlas cohorts. Figure 4A shows the total number of patients with genomic alterations in at least one of the six genes identified in our mutation network analysis (KEAP1, EP300, ERBB4, TCF12, SMC1A, NIPBL). These genes were selected based on their collective enrichment in resistant cell lines and their convergence on NFE2L2 and related cellular stress response pathways.

**Figure 4.**
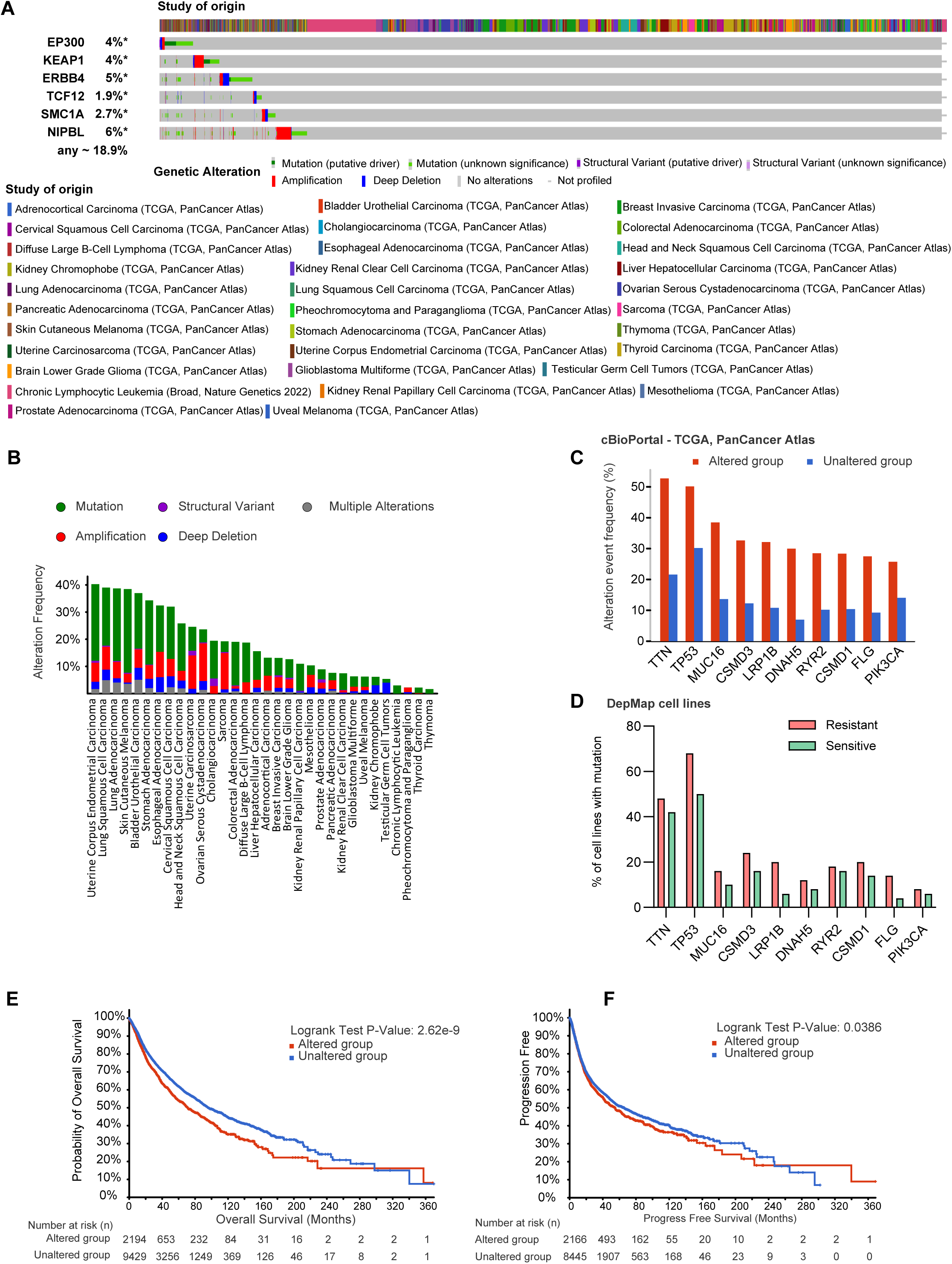
Clinical relevance of resistance-associated mutations in patient cohorts. (A) Total number of TCGA PanCancer patient samples harboring genetic alterations in at least one of six resistance-associated genes (KEAP1, EP300, ERBB4, TCF12, SMC1A, NIPBL). (B) Frequency of patients with these mutations across individual cancer types. (C) The most frequently co-occurring genomic alterations in patients with resistance-associated mutations. (D) Comparison of co-occurring mutation frequencies between resistant and sensitive DepMap cell lines. (E–F) Kaplan-Meier survival analysis (log rank test) showing reduced overall survival (E) and progression-free survival (F) in patients harboring resistance-associated mutations.

Patient samples were stratified into “altered” (with at least one genomic alteration in these genes) and “unaltered” groups, revealing broad distributions of alterations across diverse cancer types. Figure 4B presents the percentage of altered patients within each TCGA cohort, highlighting higher mutation frequencies in specific cancer types, including lung adenocarcinoma and uterine corpus endometrial carcinoma.

We next identified the most frequently co-occurring genomic alterations in the altered group (Figure 4C). Many of these co-altered genes are involved in cell signaling, transcriptional regulation, or chromatin remodeling. We then compared the mutation frequency of these co-occurring genes with our earlier analysis in sensitive and resistant lines from the DepMap dataset (Figure 4D). These alterations were more prevalent in resistant lines, further supporting their relevance to the resistance phenotype.

Finally, we analyzed clinical outcomes between altered and unaltered patient groups. Kaplan-Meier survival curves revealed significantly reduced overall survival (OS; Figure 4E) and progression-free survival (PFS; Figure 4F) in patients harboring mutations in the selected resistance-associated genes. While decreased OS likely reflects more aggressive tumor biology, the shortened PFS supports a role for these mutations in driving therapeutic resistance in human cancers.

### Drug perturbation analyses reveal re-sensitization strategies and mechanistic convergence on resistance pathways

To identify candidate compounds capable of reversing resistance phenotypes, we queried multiple drug perturbation gene set libraries using resistance-associated DEGs, prioritizing compounds predicted to repress overexpressed genes in resistant lines and restore those suppressed (Figure 5A; Supplemental Table S.13).

**Figure 5.**
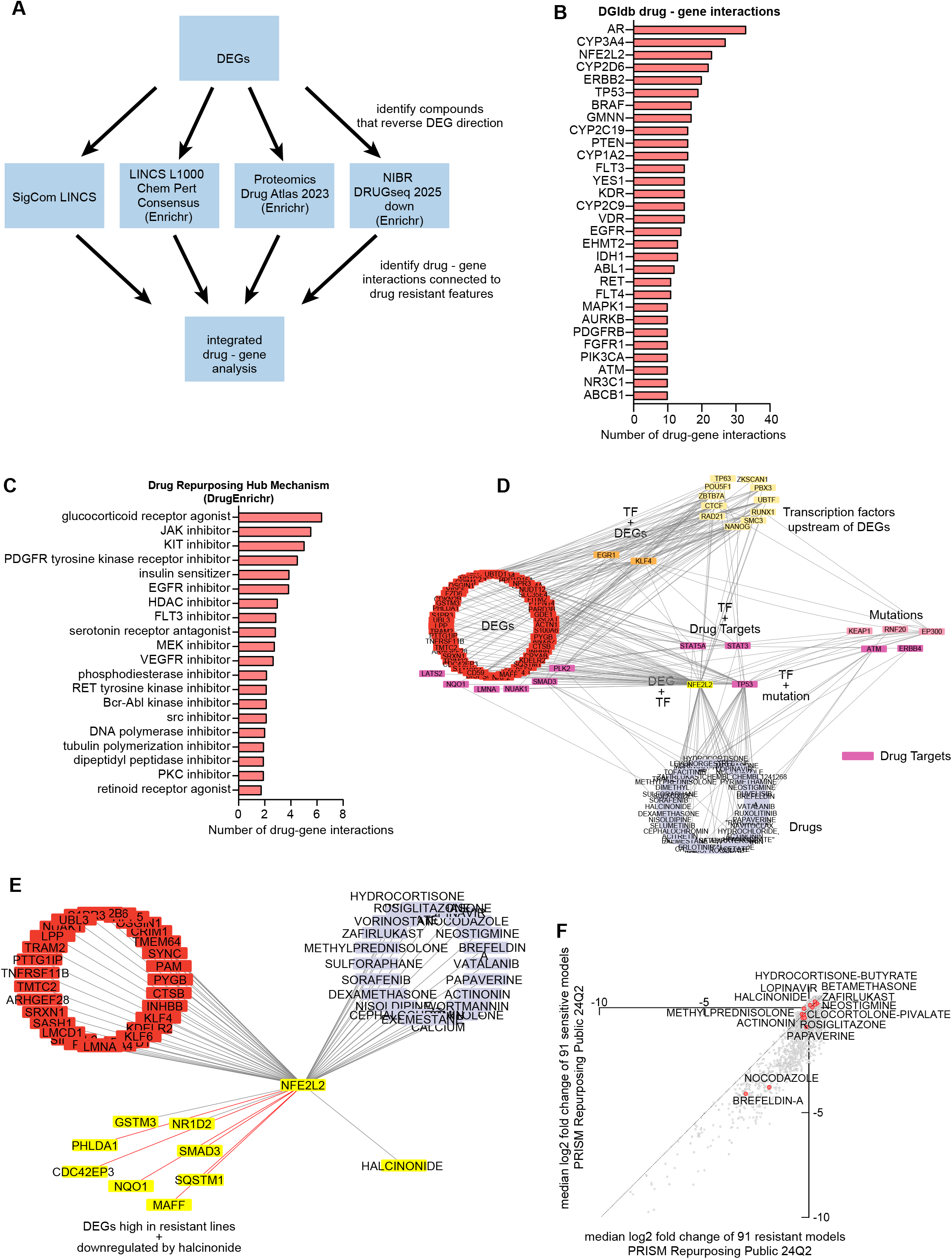
Drug perturbation analyses reveal re-sensitization strategies and mechanistic convergence on resistance pathways. (A) Schematic outlining perturbagen screening workflow using resistance-associated DEGs to identify compounds that reverse resistance-associated gene expression. (B) Top drug-gene interactions identified across datasets, highlighting recurrent targets such as NFE2L2, ABCB1, CYP3A4, and EGFR. (C) DrugEnrichR analysis of compounds for mechanisms of action. (D) Integrated drug–gene network mapping candidate compounds to resistance-associated genes and transcription factors. (E) Focused NFE2L2-centered subnetwork highlighting NFE2L2 as a target of halcinonide and DEGs predicted to be downregulated by halcinonide treatment that are also NFE2L2 targets overexpressed in resistant lines. (F) DepMap PRISM analysis of candidate compound activity, using viability as a readout (log 2 fold change) across sensitive and resistant lines, to identify bioactive compounds.

Top candidate compounds were mapped to their known gene targets using DGIdb. Several targets recurred across datasets, most notably NFE2L2, previously identified as a central transcriptional regulator in resistant lines. Other frequently targeted genes included ABCB1 (a canonical mediator of multidrug resistance through drug efflux), as well as AR, TP53, BRAF, EGFR, and various cytochrome P450 enzymes (Figure 5B). This convergence on stress response regulators (e.g., NFE2L2), drug metabolism genes (e.g., CYP3A4, CYP2D6), and efflux transporters (e.g., ABCB1) underscores a coherent mechanistic axis underlying broad resistance that may be pharmacologically reversible.

Using DrugEnrichR, we performed pathway enrichment of the candidate compounds, which revealed shared mechanisms action. The most enriched drug classes included glucocorticoid receptor agonists, JAK inhibitors, KIT and EGFR tyrosine kinase inhibitors, HDAC inhibitors, and MEK inhibitors (Figure 5C). These drug classes closely align with resistance-associated pathways identified in earlier analyses, such as stress response, chromatin remodeling, and RTK/MAPK signaling.

We then constructed a drug-gene interaction network in Cytoscape to visualize connections between resistance-targeting compounds and resistance-associated genes (Figure 5D). This network integrated DEGs and mutations from resistant lines with drug-target relationships, highlighting central hubs such as NFE2L2, EGFR, and TP53, which had also emerged in earlier transcriptomic and mutational analyses (see Figure 3G–H).

To further refine therapeutic strategies, we developed an NFE2L2-centric subnetwork highlighting its transcriptional targets upregulated in resistant lines and the compounds predicted to modulate these genes (Figure 5E). Among these, halcinonide showed strong overlap with NFE2L2 transcriptional targets and demonstrated the ability to downregulate them, supporting its potential to disrupt NFE2L2-driven resistance programs.

Finally, we leveraged DepMap PRISM repurposing data to assess the functional impact of candidate compounds in sensitive and resistant cell lines. Median log2 fold-change values were plotted for each compound in sensitive versus resistant lines (Figure 5F). Brefeldin A and nocodazole exhibited the highest overall impact on cell viability in resistant lines, suggesting these compounds effectively engage targets within resistant cells and may have the capacity to shift transcriptional programs toward a more sensitive phenotype. While not selective for resistant cells, this pharmacologic activity indicates a functional readout in resistant populations, providing a rationale for further exploration of these compounds in re-sensitization studies.

## DISCUSSION

Overcoming broad drug resistance in cancer remains a significant challenge, requiring strategies that address the multifactorial nature of therapeutic failure across diverse cancer types. Here, we developed and applied an integrative computational framework to systematically characterize broad drug resistance in cancer cell lines, connecting drug response phenotypes to multi-omic profiles, transcriptional networks, and potential re-sensitization strategies using perturbagens.

Using PRISM repurposing data from DepMap, we established a composite drug sensitivity score across mechanistically diverse compounds to stratify cell lines as broadly sensitive or resistant. This approach enabled the identification of consistent drug response phenotypes independent of individual compound or target classes, providing a foundation for subsequent omics analyses. By incorporating multiple strategies to define resistant and sensitive groups, including direct sensitivity score stratification, clustering to reduce intra-group heterogeneity, and lineage-balanced grouping, we ensured that identified features were robust and not artifacts of specific group definitions. This multi-angle approach allowed us to dissect the underlying biology of broad resistance while minimizing potential biases related to lineage or dataset-specific confounders^46-49^.

Transcriptomic analyses revealed a convergence on gene programs associated with extracellular matrix (ECM) remodeling, stress response, and survival signaling, with NFE2L2 (NRF2) emerging as a key transcriptional regulator upregulated in resistant lines. Pathway analyses further highlighted TGF-beta signaling, IGF transport, and metabolic reprogramming, including amino acid and nucleotide biosynthesis, as hallmarks of resistant phenotypes. This multi-omic convergence was reinforced by metabolomics integration, where differentially abundant metabolites and DEGs were enriched in shared pathways, underscoring the rewiring of metabolic dependencies in resistant states.

Our mutation analyses, employing both gene-level frequency comparisons and pathway-level burden analyses, revealed distinct but complementary insights. While gene-level comparisons identified pathways associated with chromatin remodeling and stress response in sensitive lines and growth factor signaling in resistant lines, pathway-level analyses across all mutations captured broader patterns of pathway enrichment, including ECM-related processes and signaling pathways relevant to resistance. Notably, the observation that Glycosaminoglycan degradation and ECM proteoglycans were enriched but directionally discordant across analysis methods emphasizes the importance of granularity in data interpretation and the complex role of ECM dynamics in modulating drug response.

Constructing an integrative gene–regulator network combining DEGs, recurrent mutations, and enriched transcription factors revealed a network of functional convergence in resistant lines. NFE2L2, a well-established regulator of oxidative stress responses and xenobiotic detoxification, previously implicated in chemoresistance across multiple cancer types^50-55^, emerged as a central hub. It connected to upstream mutations in genes such as KEAP1, EP300, ERBB4, TCF12, SMC1A, and NIPBL, and to downstream transcriptional programs involved in oxidative stress response and survival. Several of these genes, including KEAP1^56-58^ and EP300^59-61^, are well-known cancer drivers or regulators of NFE2L2 activity, and targeting this axis has recently been implicated in re-sensitization strategies^62^. Others, such as TCF12 and NIPBL, are less commonly associated with cancer or drug resistance; TCF12 is classified as a lower-confidence cancer gene in OncoKB, and NIPBL is not listed, suggesting potential novel roles in mediating transcriptional reprogramming. The tendency of these mutations to occur in a largely non-overlapping pattern while collectively impacting NFE2L2 function supports a model of convergent evolution toward a resistant phenotype through diverse genomic alterations.

Extending these findings to patient cohorts using cBioPortal and TCGA PanCancer data demonstrated that the mutation signatures identified in resistant cell lines were also prevalent in patient tumors across multiple cancer types, with strong representation in cancers known for treatment challenges, including lung and endometrial carcinomas. Importantly, patients harboring these resistance-associated mutations exhibited significantly shorter progression-free survival, supporting the translational relevance of these molecular features in mediating therapeutic resistance in human cancers.

Building on this integrative characterization, we employed resistance-associated DEGs to computationally screen perturbagen libraries, including LINCS, Drug Atlas, and DRUGseq datasets, to identify compounds capable of reversing resistance-associated transcriptional profiles. This approach leverages large-scale perturbagen datasets generated and characterized in foundational studies^24,42,44,45^, enabling systematic exploration of resistance reversal strategies. Recent advancements in perturbagen screening at high resolution, including multiplex single-cell RNA-Seq pharmacotranscriptomics^40^ and TraCe-seq clonal fitness profiling^41^, have highlighted the potential of transcriptional profiling to dissect drug response heterogeneity. While our study did not employ single-cell approaches, it complements these efforts by integrating bulk multi-omic data with perturbagen signatures to systematically prioritize candidate compounds for re-sensitization. This rational screening approach favors compounds predicted to repress genes overexpressed in resistant lines while reactivating suppressed genes, guiding drug combination selection beyond empirical methods. These findings align with broader efforts to map drug response and resistance mechanisms at scale, as exemplified by cancer dependency mapping^23^ and multi-omic functional target identification^46,63^. Drug–gene network analyses revealed convergence on stress regulators, drug metabolism genes, and efflux transporters such as NFE2L2, ABCB1, CYP3A4, and EGFR, highlighting actionable targets for re-sensitization. Focused analyses of NFE2L2-centered networks identified compounds such as halcinonide with high overlap between DEGs high in resistant lines, genes predicted to be downregulated by the perturbagen, and NFE2L2 transcriptional targets, illustrating actionable strategies for disrupting resistance pathways.

Finally, integration with DepMap PRISM drug response data demonstrated that candidate compounds such as brefeldin A and nocodazole exhibit significant activity in resistant lines, indicating target engagement and the potential to reprogram resistant cells toward a sensitive phenotype. Alternatively, compounds that reprogram resistance-associated transcriptional signatures without impacting cell viability, that could offer selective therapeutic advantages by minimizing toxicity to normal tissues, may be worth further investigating.

While this study presents a scalable computational framework for integrating multi-omics datasets with perturbagen signatures to rationally identify candidate compounds for reversing broad drug resistance, several limitations should be acknowledged. Our analyses relied on large-scale screening datasets that may not fully capture the complexity of in vivo tumor microenvironments or resistance mechanisms arising under clinical treatment pressures. Emerging single-cell perturbation approaches, such as those demonstrated in recent high-throughput pharmacotranscriptomic pipelines^40^ and TraCe-seq^41^, offer promising avenues to address these challenges by capturing intra-tumor heterogeneity and adaptive responses. Moreover, while we focused on NFE2L2 as a central regulator linking transcriptional reprogramming, stress response, and multidrug resistance, other pathways identified in our analyses warrant further exploration. For example, glycosaminoglycan metabolism and glycosylation-related pathways frequently emerged in enrichment analyses (Supplemental Tables S7–S13) but were not prioritized here. Future studies should evaluate how these pathways contribute to ECM remodeling, signaling, and resistance phenotypes. Lastly, functional validation of prioritized compounds in resistant models, including transcriptomic profiling post-treatment and preclinical testing in combinations with standard-of-care therapies, will be essential to translate these predictions into effective strategies to overcoming broad drug resistance in cancer.

## RESOURCE AVAILABILITY

### Lead Contact

Further information and requests for resources should be directed to and will be fulfilled by the lead contact, Dr. Aktar Alie.

### Materials Availability

This study did not generate new unique reagents.

### Data and Code Availability

All omics and drug screening datasets analyzed in this study are publicly available from the DepMap portal (https://depmap.org) and for the analyses of clinical samples from multiple patient cohorts in cBioPortal (https://cbioportal.org). The combined study contains samples from the following 32 studies: Acute Myeloid Leukemia (TCGA, PanCancer Atlas), Adrenocortical Carcinoma (TCGA, PanCancer Atlas), Bladder Urothelial Carcinoma (TCGA, PanCancer Atlas), Brain Lower Grade Glioma (TCGA, PanCancer Atlas), Breast Invasive Carcinoma (TCGA, PanCancer Atlas), Cervical Squamous Cell Carcinoma (TCGA, PanCancer Atlas), Cholangiocarcinoma (TCGA, PanCancer Atlas), Colorectal Adenocarcinoma (TCGA, PanCancer Atlas), Diffuse Large B-Cell Lymphoma (TCGA, PanCancer Atlas), Esophageal Adenocarcinoma (TCGA, PanCancer Atlas), Glioblastoma Multiforme (TCGA, PanCancer Atlas), Head and Neck Squamous Cell Carcinoma (TCGA, PanCancer Atlas), Kidney Chromophobe (TCGA, PanCancer Atlas), Kidney Renal Clear Cell Carcinoma (TCGA, PanCancer Atlas), Kidney Renal Papillary Cell Carcinoma (TCGA, PanCancer Atlas), Liver Hepatocellular Carcinoma (TCGA, PanCancer Atlas), Lung Adenocarcinoma (TCGA, PanCancer Atlas), Lung Squamous Cell Carcinoma (TCGA, PanCancer Atlas), Mesothelioma (TCGA, PanCancer Atlas), Ovarian Serous Cystadenocarcinoma (TCGA, PanCancer Atlas), Pancreatic Adenocarcinoma (TCGA, PanCancer Atlas), Pheochromocytoma and Paraganglioma (TCGA, PanCancer Atlas), Prostate Adenocarcinoma (TCGA, PanCancer Atlas), Sarcoma (TCGA, PanCancer Atlas), Skin Cutaneous Melanoma (TCGA, PanCancer Atlas), Stomach Adenocarcinoma (TCGA, PanCancer Atlas), Testicular Germ Cell Tumors (TCGA, PanCancer Atlas), Thymoma (TCGA, PanCancer Atlas), Thyroid Carcinoma (TCGA, PanCancer Atlas), Uterine Carcinosarcoma (TCGA, PanCancer Atlas), Uterine Corpus Endometrial Carcinoma (TCGA, PanCancer Atlas), Uveal Melanoma (TCGA, PanCancer Atlas).

Processed data and custom analyses performed using DepMap portal used to generate figures and perform pathway/network analyses available in supplemental tables.

Additional supporting data are available upon request from the lead contact.

## ACKNOWLEDGMENTS

This work was supported by funding from the Warren Family Research Center for Drug Discovery and Development. We also acknowledge the University of Notre Dame Biological Screening and Development Core for resources.

## AUTHOR CONTRIBUTIONS

I.M.: Conceptualization, methodology, investigation, formal analysis, data curation, visualization, writing – original draft. B.B.: Supervision, project administration, funding acquisition. A.A.: Conceptualization, supervision, funding acquisition, writing – review & editing.

## DECLARATION OF INTERESTS

The authors declare no competing interests.

### Declaration of Generative AI and AI-Assisted Technologies in the Writing Process

During the preparation of this work, the authors used OpenAI’s ChatGPT to assist with editing, and language refinement. After using this tool, the authors reviewed and edited the content as needed and take full responsibility for the content of the publication.

## SUPPLEMENTAL INFORMATION

Document S1 (Supplemental material related to Figure 2)

Tables S1-S13 (Excel file with Supplemental Tables S1 – S13)

## METHODS

### Drug Sensitivity Data and Calculation of Global Resistance Scores

Drug sensitivity data were obtained from the PRISM Repurposing dataset (DepMap 24Q1), containing log_2_ fold-change (LFC) viability scores across 6765 compounds in cancer cell lines. For each cell line, median LFC across all compounds and class-specific medians (kinase inhibitors, chemotherapies, epigenetic modulators) were calculated. A composite global drug sensitivity score was computed by averaging class-level medians to stratify cell lines as broadly resistant or sensitive.

### Selection and Refinement of Resistant and Sensitive Cell Lines

Cell lines were grouped into sensitive and resistant populations using 3 different strategies. (1) Cell lines in the top and bottom deciles of drug sensitivity scores were classified as resistant or sensitive, respectively. (2) To reduce intra-group heterogeneity, a supervised feature selection was performed using correlation analysis (expression vs. sensitivity score) followed by ANOVA F-test to retain the top 250 discriminatory genes. Z-score normalization was applied, and hierarchical clustering (Ward’s method, Euclidean distance) was performed to identify the largest contiguous, phenotype-pure clusters for downstream analyses. (3) Lineage balanced groups were constructed to control for lineage-specific biases by including only cancer types represented in both resistant and sensitive categories (top 10%) and selecting an equal number of a specific lineage based on highest or lowest drug sensitivity score for resistant and sensitive lines respectively.

### Metadata Association Analyses

Metadata from DepMap (lineage, disease subtype, collection site, media, age) were tested for associations with drug sensitivity scores using univariate linear regression (OLS) models to assess potential confounders.

### Differential Gene Expression and Metabolomics Analysis

Differential gene expression analyses were performed using the DepMap Custom Analysis tool (https://depmap.org/portal/interactive/custom_analysis), conducting two-class comparisons between resistant and sensitive cell line subsets defined by composite drug sensitivity scores. This tool computes effect sizes (interpreted as fold differences between groups) and P-values using its internal pipelines on RNA-seq gene expression data reported as log_2_(CPM + 1) for each gene across cell lines. Genes with P < 0.05 and effect size > 1 were considered significant and an additional filtering criteria was used, which included a correlation analysis between drug sensitivity score and gene expression (log_2_(CPM+1) values from DepMap). Genes shared across our 3 separate for downstream analyses. Differential metabolite abundance analyses were also conducted using the DepMap Custom Analysis tool, performing two-class comparisons on resistant and sensitive groups.

### Pathway and Transcription Factor Enrichment Analysis

Pathway enrichment analyses were performed using Enrichr^64^ (https://maayanlab.cloud/Enrichr/), querying Reactome and ENCODE/ChEA consensus transcription factor databases on upregulated and downregulated DEGs.

### Metabolomics Data Integration and Joint Pathway Analysis

Metabolomics data were integrated using differentially abundant metabolites from DepMap. Combined DEG and metabolite sets were analyzed in MetaboAnalyst 5.0^65^ (https://www.metaboanalyst.ca).

### Mutation Analysis and Pathway-Level Burden Scoring

Mutation data (binary gene-level mutation presence) were retrieved from DepMap. Gene-level mutation frequency skew scores (resistant vs. sensitive) were computed, and the top 50 skewed genes were subjected to pathway enrichment via Enrichr.

Sample-wise pathway mutation burden scoring was performed using Reactome pathway gene sets, computing – log_10_(p-value) enrichment scores on each individual sample and generating heatmaps and volcano plots summarizing pathway-level enrichment and group skew. Group skew was defined as the difference in the % of lines in a population with significant enrichment to a pathway, and a correction factor using -log_10_(p-value).

### Network Construction and Visualization

Gene–regulator networks were constructed by integrating DEGs, enriched transcription factors (ENCODE/ChEA), and frequently mutated genes using Cytoscape 3.9.1^66^ (https://cytoscape.org) for visualization. Focused subnetworks centered on NFE2L2 and associated modules were generated for pathway-centric analyses.

### Patient Cohort Analyses in cBioPortal

Clinical relevance was assessed using cBioPortal^67^ (https://www.cbioportal.org), querying TCGA PanCancer Atlas^68^ data for mutations in KEAP1, EP300, ERBB4, TCF12, SMC1A, and NIPBL across 32 cancer types. Mutation frequencies, co-occurrence analyses, and Kaplan-Meier survival plots (log-rank test) for overall and progression-free survival were generated using integrated cBioPortal tools.

### Perturbagen Screening and Drug–Gene Network Analyses

Resistance-associated DEGs were used to query LINCS L1000, Drug Atlas, and DRUGseq perturbation libraries to identify compounds predicted to reverse resistant gene expression signatures, leveraging SigCom LINCS^42^ (https://maayanlab.cloud/sigcom-lincs) and Connectivity Map resources. Enrichment for drug mechanisms of action was performed using DrugEnrichr^69^ (https://maayanlab.cloud/DrugEnrichr). Drug–gene interactions were mapped using DGIdb 4.2^70^ (https://www.dgidb.org), and networks were visualized in Cytoscape to identify convergence on NFE2L2.

### Compound Activity Validation Using DepMap PRISM

Candidate compound activities were evaluated by extracting PRISM repurposing dataset viability data (log_2_ fold change) in resistant and sensitive lines to confirm bioactivity profiles and prioritize compounds for re-sensitization potential.

---

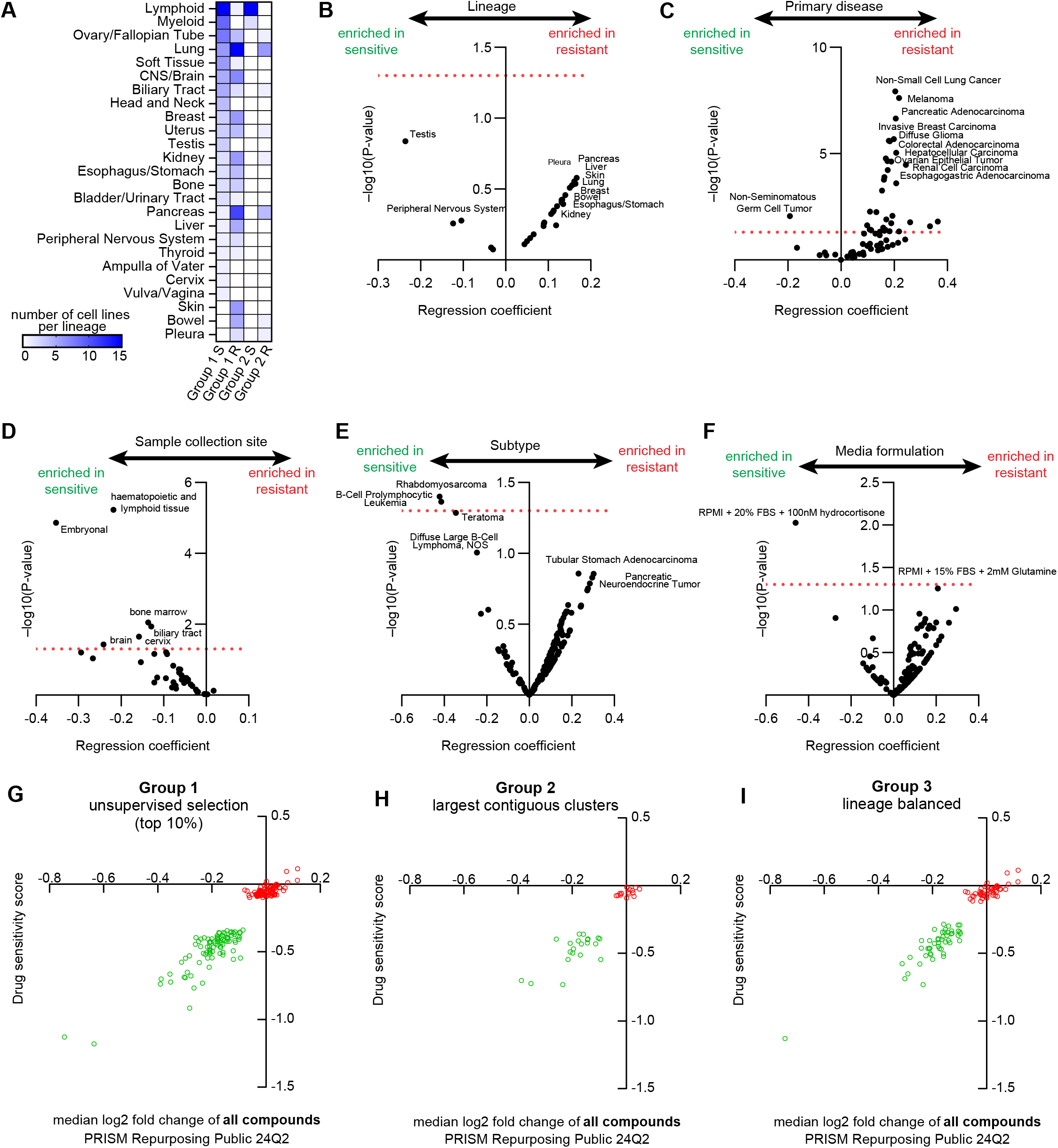

## References

1 Baguley, B. C. Multiple drug resistance mechanisms in cancer. Molecular biotechnology 46, 308–316 (2010).

2 Szakács, G., Paterson, J. K., Ludwig, J. A., Booth-Genthe, C. & Gottesman, M. M. Targeting multidrug resistance in cancer. Nature reviews Drug discovery 5, 219–234 (2006).

3 Ozben, T. Mechanisms and strategies to overcome multiple drug resistance in cancer. FEBS letters 580, 2903–2909 (2006).

4 Fletcher, J. I., Williams, R. T., Henderson, M. J., Norris, M. D. & Haber, M. ABC transporters as mediators of drug resistance and contributors to cancer cell biology. Drug Resistance Updates 26, 1–9 (2016).

5 Robey, R. W. et al. Revisiting the role of efflux pumps in multidrug-resistant cancer. Nature reviews. Cancer 18, 452 (2018).

6 Krishna Vadlapatla, R., Dutt Vadlapudi, A., Pal, D. & K Mitra, A. Mechanisms of drug resistance in cancer chemotherapy: coordinated role and regulation of efflux transporters and metabolizing enzymes. Current pharmaceutical design 19, 7126–7140 (2013).

7 Assaraf, Y. G. The role of multidrug resistance efflux transporters in antifolate resistance and folate homeostasis. Drug Resistance Updates 9, 227–246 (2006).

8 Kim, P., Li, H., Wang, J. & Zhao, Z. Landscape of drug-resistance mutations in kinase regulatory hotspots. Briefings in bioinformatics 22, bbaa108 (2021).

9 Brown, J. M. & Attardi, L. D. The role of apoptosis in cancer development and treatment response. Nature reviews cancer 5, 231–237 (2005).

10 Schmitt, C. A. & Lowe, S. W. Apoptosis is critical for drug response in vivo. Drug Resistance Updates 4, 132–134 (2001).

11 Borst, P., Borst, J. & Smets, L. A. Does resistance to apoptosis affect clinical response to antitumor drugs? Drug resistance updates: reviews and commentaries in antimicrobial and anticancer chemotherapy 4, 129–131 (2001).

12 Lines, C. L., McGrath, M. J., Dorwart, T. & Conn, C. S. The integrated stress response in cancer progression: a force for plasticity and resistance. Frontiers in Oncology 13, 1206561 (2023).

13 Pazarentzos, E. & Bivona, T. G. Adaptive stress signaling in targeted cancer therapy resistance. Oncogene 34, 5599–5606 (2015).

14 Tomida, A. & Tsuruo, T. Drug resistance mediated by cellular stress response to the microenvironment of solid tumors. Anti-cancer drug design 14, 169–177 (1999).

15 Borst, P. Cancer drug pan-resistance: pumps, cancer stem cells, quiescence, epithelial to mesenchymal transition, blocked cell death pathways, persisters or what? Open biology 2, 120066 (2012).

16 Liu, P. et al. Disulfiram targets cancer stem-like cells and reverses resistance and cross-resistance in acquired paclitaxel-resistant triple-negative breast cancer cells. British journal of cancer 109, 1876–1885 (2013).

17 Phi, L. T. H. et al. Cancer stem cells (CSCs) in drug resistance and their therapeutic implications in cancer treatment. Stem cells international 2018, 5416923 (2018).

18 Donnenberg, V. S. & Donnenberg, A. D. Multiple drug resistance in cancer revisited: the cancer stem cell hypothesis. The Journal of Clinical Pharmacology 45, 872–877 (2005).

19 Prieto-Vila, M., Takahashi, R.-u., Usuba, W., Kohama, I. & Ochiya, T. Drug resistance driven by cancer stem cells and their niche. International journal of molecular sciences 18, 2574 (2017).

20 Dubikovskaya, E. A., Thorne, S. H., Pillow, T. H., Contag, C. H. & Wender, P. A. Overcoming multidrug resistance of small-molecule therapeutics through conjugation with releasable octaarginine transporters. Proceedings of the National Academy of Sciences 105, 12128–12133 (2008).

21 Shao, N. et al. Reversing anticancer drug resistance by synergistic combination of chemotherapeutics and membranolytic antitumor β-peptide polymer. Journal of the American Chemical Society 146, 11254–11265 (2024).

22 Gottesman, M. M., Robey, R. W. & Ambudkar, S. V. New mechanisms of multidrug resistance: an introduction to the Cancer Drug Resistance special collection. Cancer Drug Resistance 6, 590 (2023).

23 Arafeh, R., Shibue, T., Dempster, J. M., Hahn, W. C. & Vazquez, F. The present and future of the Cancer Dependency Map. Nature Reviews Cancer 25, 59–73 (2025). 10.1038/s41568-024-00763-x

24 Corsello, S. M. et al. Discovering the anticancer potential of non-oncology drugs by systematic viability profiling. Nature Cancer 1, 235–248 (2020). 10.1038/s43018-019-0018-6

25 Bondeson, D. P. et al. Phosphate dysregulation via the XPR1–KIDINS220 protein complex is a therapeutic vulnerability in ovarian cancer. Nature cancer 3, 681–695 (2022).

26 Ishizuka, J. J. et al. Loss of ADAR1 in tumours overcomes resistance to immune checkpoint blockade. Nature 565, 43–48 (2019).

27 Manguso, R. T. et al. In vivo CRISPR screening identifies Ptpn2 as a cancer immunotherapy target. Nature 547, 413–418 (2017).

28 Dixit, A. et al. Perturb-Seq: dissecting molecular circuits with scalable single-cell RNA profiling of pooled genetic screens. cell 167, 1853-1866. e1817 (2016).

29 Oprea, T. I. et al. Unexplored therapeutic opportunities in the human genome. Nature reviews Drug discovery 17, 317–332 (2018).

30 Tanoli, Z. et al. Computational drug repurposing: approaches, evaluation of in silico resources and case studies. Nature Reviews Drug Discovery (2025). 10.1038/s41573-025-01164-x

31 Finan, C. et al. The druggable genome and support for target identification and validation in drug development. Science translational medicine 9, eaag1166 (2017).

32 Piñero, J. et al. DisGeNET: a discovery platform for the dynamical exploration of human diseases and their genes. Database 2015, bav028 (2015).

33 McDonagh, E. M. et al. Human genetics and genomics for drug target identification and prioritization: Open Targets’ perspective. Annual review of biomedical data science 7 (2024).

34 Sondka, Z. et al. The COSMIC Cancer Gene Census: describing genetic dysfunction across all human cancers. Nature Reviews Cancer 18, 696–705 (2018).

35 Tomczak, K., Czerwińska, P. & Wiznerowicz, M. Review The Cancer Genome Atlas (TCGA): an immeasurable source of knowledge. Contemporary Oncology/Współczesna Onkologia 2015, 68–77 (2015).

36 Landrum, M. J. et al. ClinVar: public archive of relationships among sequence variation and human phenotype. Nucleic acids research 42, D980–D985 (2014).

37 Huang, Y.-Q. et al. Bioinformatics toolbox for exploring target mutation-induced drug resistance. Briefings in Bioinformatics 24 (2023). 10.1093/bib/bbad033

38 Wang, L. et al. Unbiased discovery of cancer pathways and therapeutics using Pathway Ensemble Tool and Benchmark. Nature Communications 15, 7288 (2024). 10.1038/s41467-024-51859-9

39 Aissa, A. F. et al. Single-cell transcriptional changes associated with drug tolerance and response to combination therapies in cancer. Nature Communications 12, 1628 (2021). 10.1038/s41467-021-21884-z

40 Dini, A. et al. A multiplex single-cell RNA-Seq pharmacotranscriptomics pipeline for drug discovery. Nature Chemical Biology 21, 432–442 (2025). 10.1038/s41589-024-01761-8

41 Chang, M. T. et al. Identifying transcriptional programs underlying cancer drug response with TraCe-seq. Nature Biotechnology 40, 86–93 (2022). 10.1038/s41587-021-01005-3

42 Evangelista, J. E. et al. SigCom LINCS: data and metadata search engine for a million gene expression signatures. Nucleic Acids Research 50, W697–W709 (2022). 10.1093/nar/gkac328

43 Rees, M. G. et al. Correlating chemical sensitivity and basal gene expression reveals mechanism of action. Nature Chemical Biology 12, 109–116 (2016). 10.1038/nchembio.1986

44 Gonçalves, E. et al. Drug mechanism-of-action discovery through the integration of pharmacological and CRISPR screens. Molecular Systems Biology 16, e9405 (2020).

45 Subramanian, A. et al. A next generation connectivity map: L1000 platform and the first 1,000,000 profiles. Cell 171, 1437-1452. e1417 (2017).

46 Warren, A. et al. Global computational alignment of tumor and cell line transcriptional profiles. Nature communications 12, 22 (2021).

47 Noorbakhsh, J., Vazquez, F. & McFarland, J. M. Bridging the gap between cancer cell line models and tumours using gene expression data. British Journal of Cancer 125, 311–312 (2021).

48 Corsello, S. M. et al. Discovering the anticancer potential of non-oncology drugs by systematic viability profiling. Nature cancer 1, 235–248 (2020).

49 Barretina, J. et al. The Cancer Cell Line Encyclopedia enables predictive modelling of anticancer drug sensitivity. Nature 483, 603–607 (2012).

50 Liu, X., Xu, C., Xiao, W. & Yan, N. Unravelling the role of NFE2L1 in stress responses and related diseases. Redox biology 65, 102819 (2023).

51 Liu, Z. et al. Association of KEAP1 and NFE2L2 polymorphisms with temporal lobe epilepsy and drug resistant epilepsy. Gene 571, 231–236 (2015).

52 Dempke, W. C. & Reck, M. KEAP1/NRF2 (NFE2L2) mutations in NSCLC–Fuel for a superresistant phenotype? Lung cancer 159, 10–17 (2021).

53 Hallis, S. P., Go, B. J., Yoo, J. M., Cho, G. H. & Kwak, M.-K. Toward a Better Understanding of NRF2/NFE2L2 and BCRP/ABCG2 in Therapy Resistance in Cancer. Drug Targets and Therapeutics 2, 111–123 (2023).

54 Maiti, A. K. Overcoming drug resistance through elevation of ROS in cancer. Molecular Mechanisms of Tumor Cell Resistance to Chemotherapy: Targeted Therapies to Reverse Resistance, 135-149 (2013).

55 Rajesh, Y. et al. Targeting NFE2L2, a transcription factor upstream of MMP-2: a potential therapeutic strategy for temozolomide resistant glioblastoma. Biochemical Pharmacology 164, 1–16 (2019).

56 Kansanen, E., Kuosmanen, S. M., Leinonen, H. & Levonen, A.-L. The Keap1-Nrf2 pathway: Mechanisms of activation and dysregulation in cancer. Redox biology 1, 45–49 (2013).

57 Taguchi, K. & Yamamoto, M. The KEAP1–NRF2 system as a molecular target of cancer treatment. Cancers 13, 46 (2020).

58 Taguchi, K. & Yamamoto, M. The KEAP1–NRF2 system in cancer. Frontiers in oncology 7, 85 (2017).

59 Karunatilleke, N. C., Brickenden, A. & Choy, W. Y. Molecular basis of the interactions between the disordered Neh4 and Neh5 domains of Nrf2 and CBP/p300 in oxidative stress response. Protein Science 33, e5137 (2024).

60 Vo, N. & Goodman, R. H. CREB-binding protein and p300 in transcriptional regulation. Journal of Biological Chemistry 276, 13505–13508 (2001).

61 Gronkowska, K. & Robaszkiewicz, A. Genetic dysregulation of EP300 in cancers in light of cancer epigenome control–targeting of p300-proficient and-deficient cancers. Molecular Therapy Oncology 32 (2024).

62 Wang, K., Baird, L. & Yamamoto, M. The clinical-grade CBP/p300 inhibitor CCS1477 represses the global NRF2-dependent cytoprotective transcription program and re-sensitizes cancer cells to chemotherapeutic drugs. Free Radical Biology and Medicine 233, 102–117 (2025).

63 Pacini, C. et al. Integrated cross-study datasets of genetic dependencies in cancer. Nature Communications 12, 1661 (2021). 10.1038/s41467-021-21898-7

64 Chen, E. Y. et al. Enrichr: interactive and collaborative HTML5 gene list enrichment analysis tool. BMC Bioinformatics 14, 128 (2013). 10.1186/1471-2105-14-128

65 Pang, Z. et al. MetaboAnalyst 5.0: narrowing the gap between raw spectra and functional insights. Nucleic Acids Research 49, W388–W396 (2021). 10.1093/nar/gkab382

66 Shannon, P. et al. Cytoscape: a software environment for integrated models of biomolecular interaction networks. Genome research 13, 2498–2504 (2003).

67 Gao, J. et al. Integrative analysis of complex cancer genomics and clinical profiles using the cBioPortal. Science signaling 6, pl1–pl1 (2013).

68 Weinstein, J. N. et al. The cancer genome atlas pan-cancer analysis project. Nature genetics 45, 1113–1120 (2013).

69 Kuleshov, M. V. et al. modEnrichr: a suite of gene set enrichment analysis tools for model organisms. Nucleic Acids Research 47, W183–W190 (2019). 10.1093/nar/gkz347

70 Cannon, M. et al. DGIdb 5.0: rebuilding the drug–gene interaction database for precision medicine and drug discovery platforms. Nucleic acids research 52, D1227–D1235 (2024).

